# A novel PD-L1-targeted shark V_NAR_ single domain-based CAR-T strategy for treating breast cancer and liver cancer

**DOI:** 10.1101/2021.07.20.453144

**Authors:** Dan Li, Hejiao English, Jessica Hong, Tianyuzhou Liang, Glenn Merlino, Chi-Ping Day, Mitchell Ho

**Affiliations:** Laboratory of Molecular Biology, Center for Cancer Research, National Cancer Institute, Bethesda, MD; Laboratory of Cancer Biology and Genetics, Center for Cancer Research, National Cancer Institute, Bethesda, MD

**Author notes:** The work was done in Bethesda, MD, USA. Correspondence should be addressed to Mitchell Ho, Mitchell Ho, Ph.D., Laboratory of Molecular Biology, National Cancer Institute, National Institutes of Health, 37 Convent Drive, Room 5002, Bethesda, MD 20892-4264, USA.

**Keywords:** shark V_NAR_, single domain antibody, CAR-T cells, immune checkpoint, PD-L1, triple-negative breast cancer, hepatocellular carcinoma or HCC, liver cancer, glypican-3 or GPC3, xenograft

## Abstract

Chimeric antigen receptor (CAR)-T cell therapy shows great potency against hematological malignancies, whereas it remains difficult to treat solid tumors mainly due to lack of appropriate antigenic targets and immunosuppressive tumor microenvironment (TME). Checkpoint molecule PD-L1 is widely overexpressed on multiple tumor types, and the PD-1/PD-L1 interaction is a key mediator of immunosuppression in TME. Here, we isolated anti-PD-L1 single domain antibodies from a newly constructed semi-synthetic nurse shark V_NAR_ phage library. We found that one V_NAR_, B2, showed cross-reactivity to human, mouse, and canine PD-L1 antigens, and it partially blocked the interaction of human PD-1 to PD-L1. Furthermore, CAR (B2) T cells specifically lysed human breast cancer and liver cancer cells by targeting constitutive and inducible expression of PD-L1, and also hindered tumor metastasis. Importantly, the combination of CAR (B2) T cells with CAR-T cells targeting liver cancer-specific antigen GPC3 regress liver tumors in mice. We concluded that PD-L1-targeted shark V_NAR_ single domain-based CAR-T therapy is a novel strategy to treat breast cancer and liver cancer. This provides a rationale for potential use of CAR (B2) T cells as a monotherapy or combination with a tumor-specific therapy in clinical studies.

## Background

Adoptive cell therapy (ACT), particularly chimeric antigen receptor (CAR)-T cell therapy, has shown great potency as one of the most effective cancer immunotherapies^1–3^. CARs are synthetic receptors consisting of an extracellular domain, a hinge region, a transmembrane domain, and intracellular signal domains (e.g. CD3-zeta, CD28, 41BB) that initiate T cell activation^4–6^. CARs can promote non-major histocompatibility complex (MHC)-restricted recognition of cell surface components, bind tumor antigens directly, and trigger a T-cell anti-tumor response^7^. CAR-T cells targeting B cell antigen CD19 have shown clinical success in patients with advanced B cell lymphoma, which led to their approval by the U.S. Food and Drug Administration (FDA)^3,8^. However, the translation of CAR-T cells to solid tumors is more difficult because of a lack of appropriate antigenic targets and the complex immunosuppressive tumor microenvironment (TME). Recently, the proteins glypican-2 (GPC2)^9^, glypican-3 (GPC3)^10^, and mesothelin^11,12^ were reported as emerging antigens for CAR-T therapy in the treatment of solid tumors and development for clinical trials. However, not all tumors express highly specific surface antigens that are suitable for CARs recognition. Tumor heterogeneity makes targeted therapy more challenging. Programmed death-ligand 1 (PD-L1 or CD274) has aberrantly high expression on multiple tumor types through oncogenic signaling^13^, and is induced by pro-inflammatory factors such as IFN-γ in the immune-reactive TME^14^. It has been shown that PD-L1 expressed on tumors can induce T-cell tolerance and avoid immune destruction through binding with its ligand programmed cell death protein 1 (PD-1) on T-cell, which may be one of the main reasons for the poor effect of CAR-T in solid tumors^15^. Clinically, antibody-based PD-1/PD-L1 antagonists were reported to induce durable tumor inhibition, especially in melanoma, non-small cell lung cancer, and renal cancer. However, the response rate remains poor in other types of advanced solid tumor^16^. Recently, PD-L1-targeting camelid V_H_H-nanobody-based CAR-T cells have shown to delay tumor growth in a syngeneic mouse melanoma model^17^. Moreover, PD-L1-targeting CAR natural killer (NK) cells inhibited the growth of triple-negative breast cancer (TNBC), lung cancer, and bladder tumors engrafted in NOD SCID gamma (NSG) mice^18^. Furthermore, bi-specific Trop2/PD-L1 CAR-T cells targeting both Trop2 and PD-L1 demonstrated improved killing effect of CAR-T cells in gastric cancer^19^. PD-L1-targeted CAR-T cell therapy is presumed to kill PD-L1-overexpressing tumor cells and block the PD-1/PD-L1 immune checkpoint, thereby significantly enhancing anti-tumor activity in solid tumors.

The single-chain antibody variable fragment (scFv) commonly serves as the antigen-recognition region of a CAR construct, which consists of heavy (V_H_) and variable light (V_L_) chains connected by a flexible linker (Gly_4_Ser)_3_. However, folding of an artificially engineered scFv can affect the specificity and affinity of the CAR for its target antigen^20^. In contrast, the antigen binding domain of naturally occurring single-domain antibodies (heavy chain-only) from camelid (V_H_H)^21^ and shark (V_NAR_)^22^ have beneficial properties for the engineering of CARs. They are small in size (12-15 kDa), easily expressed, and capable of binding concave and hidden epitopes that are not accessible to conventional antibodies^23^. Remarkably, shark V_NAR_s have unique features that are distinct from camel V_H_Hs—they are in large diversity, and are evolutionally derived from an ancient single domain that functions as a variable domain in both B cell and T cell receptors^24,25^. We previously constructed a V_NAR_ phage-displayed library from six nurse sharks^26^. Currently, there are several shark V_NAR_s emerging from pre-clinical research. Their therapeutic and biotechnological applications are under intensive investigation^27–29^.

In this study, we reconstructed a semi-synthetic shark V_NAR_ phage library with randomized third complementarity-determining regions (CDR3) of 18 amino acids (AA) in length. Of the three binders that were cross-reactive with mouse and human antigens, only B2 could functionally block the interaction between human PD-L1 and PD-1. More importantly, B2-based CAR-T cells successfully inhibited tumor growth in the xenograft mouse models of TNBC and hepatocellular carcinoma (HCC). Interestingly, the combination of CAR (B2) T cells and liver cancer specific GPC3 CAR demonstrated better efficacy in a synergistic manner compared to single antigen-targeted CAR-T cells in mice, highlighting the feasibility and efficacy of PD-L1-targeting shark V_NAR_-CAR-T cells in solid tumors.

## Results

### Construction of a semi-synthetic shark V_NAR_ single domain library

We previously constructed a naïve shark V_NAR_ library from 6 naïve adult nurse sharks (*Ginglymostoma cirratum*) with a size of 1.2 × 10^10^ pfu/ml ^25,26^. To improve the diversity and utility of the shark V_NAR_ library, in this study we developed a semi-synthetic randomized CDR3 18AA shark V_NAR_ library (referred to as ‘18AA CDR3 shark library’). As illustrated in Fig. 1A, 70% of V_NAR_s in the naïve nurse shark library are type II, containing two canonical cysteines located at amino acid 21 and 82 to form a disulfide bond and at least one extra cysteine in CDR1 and CDR3 to form an interloop disulfide bond. Since the type IV V_NAR_ sequence is the closest to its mammalian counterpart such as human V_H_ with only a pair of canonical cysteines, one before CDR1 and the other before CDR3, we made the C29Y mutation and randomized CDR3 loop region to change all V_NAR_s to type IV instead of four types (type I, II, III, and IV). The diversity of the newly semi-synthetic library is 1.2 × 10^10^ pfu/ml which is comparable with the naïve shark V_NAR_ library (Fig. 1A and 1B). To assess the randomness of sequence modification, we estimated the average nucleotide ratio at each CDR3 residue based on sequencing analysis and found that the CDR3 nucleotides were completely randomized with desired ATGC bases ratios (Fig. 1C).

**Figure 1.**
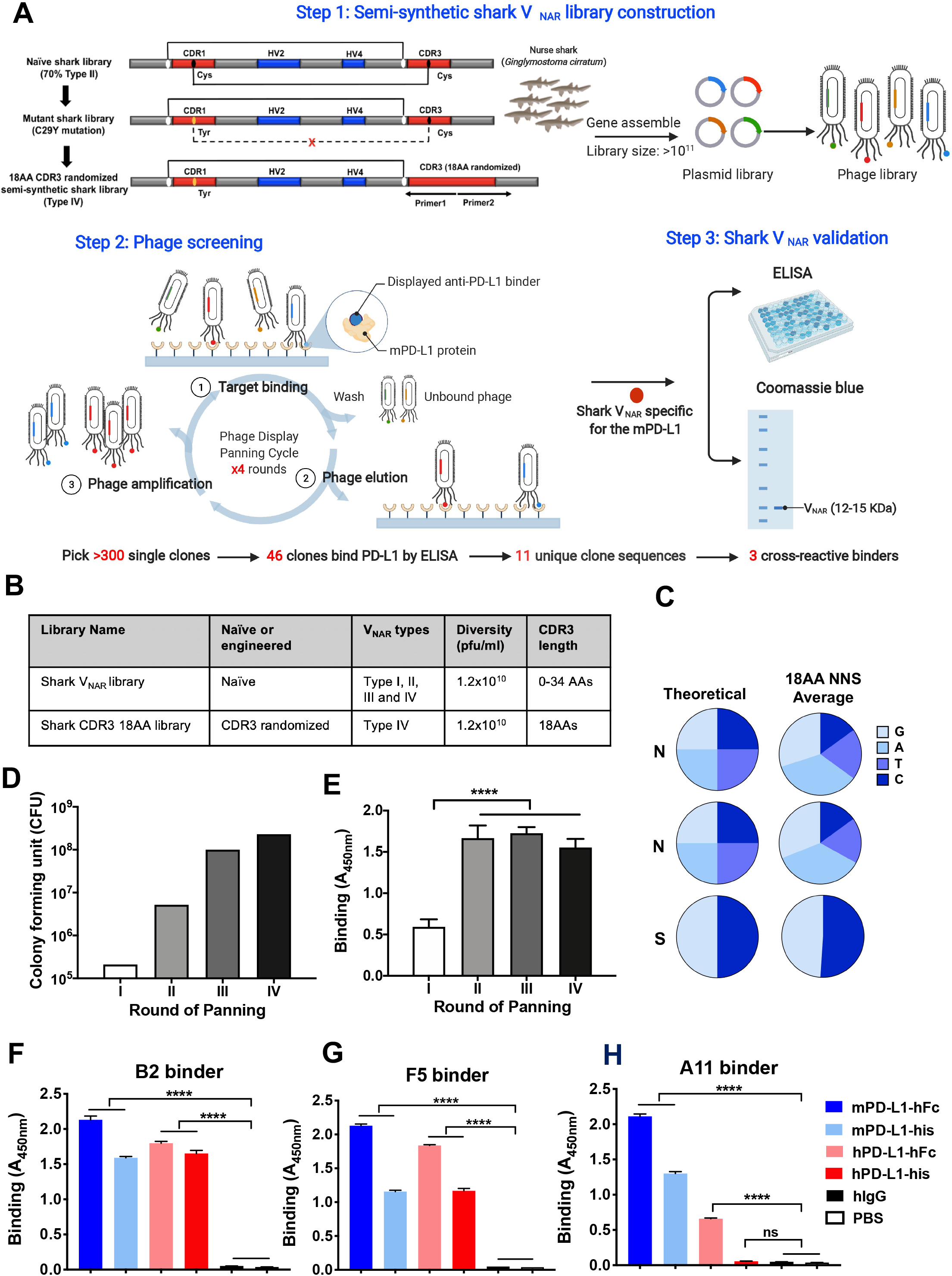
Isolation of anti-PD-L1 single domain antibody by phage display from an engineered semi-synthetic shark V_NAR_ phage library. (A) Circuit of three steps library construction and phage panning. A 18AA randomized CDR3 semi-synthetic shark V_NAR_ phage library was constructed by PCR mutation and gene assemble. After 3-5 rounds of phage panning, anti-mPD-L1 V_NAR_s were isolated from the phage library, and further validated by phage ELISA and protein purification technologies. (B) Information regarding newly shark V_NAR_ library compared with pre-synthetic V_NAR_ library. (C) Pie chart of the percentage of average nucleotide (ACTG) ratio at each randomization NNS. (D) Phage-displayed single-domain antibody clones were identified against recombinant mPD-L1-his after four rounds of panning. A gradual increase in phage titers was observed during each round of panning. (E) Polyclonal phage ELISA from the output phage of each round of panning. (F-H) Cross-reactivity of anti-PD-L1 B2 (F), A11 (G), and F5 (H) to mPD-L1 and hPD-L1 protein within His-tag or hFc-tag by monoclonal phage ELISA analysis.

### Isolation of cross-species V_NAR_ single domains with high affinity for PD-L1

To identify the anti-PD-L1 shark V_NAR_ that can play a role in the murine tumor environment, we used mouse PD-L1 (mPD-L1) protein as an antigen to screen the new semi-synthetic shark library (Fig. 1A). After four rounds of panning, ≈1,000-fold enrichment of eluted phage colonies was obtained (Fig. 1D). We also observed an enhanced binding to PD-L1 after the first round of phage panning (Fig. 1E). At the end of the fourth round of panning, 46 individual clones were identified to bind mPD-L1 protein by the monoclonal phage enzyme-linked immunosorbent assay (ELISA), and 11 unique binders were confirmed by subsequent sequencing. Three PD-L1-specific V_NAR_s, B2, A11, and F5, finally showed cross-reactivity to both mouse (mPD-L1) and human PD-L1 (hPD-L1) protein in either His-tag or the hFc-tag formats, as shown by monoclonal phage ELISA (Fig. 1F-H).

To determine the antigen specificity of shark V_NAR_s, we established three PD-L1 knockout (KO) single clones by the CRISPR-Cas9 technology in a human TNBC cell line, MDA-MB-231. To enhance the PD-L1 knockout efficiency, two single guide RNAs (sgRNAs) were designed to target the promoter of the endogenous PD-L1 gene (Fig. 2A). All three individual cell clones confirmed the loss of PD-L1 expression (Fig. 2A), and clone 1 was further used in the present study. To determine cross-species reactivity of anti-PD-L1 shark V_NAR_s against native PD-L1, three PD-L1 positive tumor cell lines, including a human breast cancer cell line, a mouse melanoma cell line, and a canine melanoma cell line, were used to evaluate binding activity of B2, A11, and F5. As shown in Fig. 2B, both B2 and F5 bind human antigen, and cross-react with mouse and canine antigens. B2 showed a higher binding ability to both human and mouse antigens than that of F5. A11 bind canine antigen but not antigen of human or mouse. In contrast, no binding was shown on PD-L1 KO cells, indicating the binding activity of shark V_NAR_s is antigen-specific. To determine binding kinetics, we further produced V_NAR_-Fc fusion protein and incubated them with hPD-L1-His protein on the Bio-layer interferometry (BLI) Octet platform. The K_D_ value of the B2 was 1.7 nM and 1.4 nM at a concentration of 100 nM and 50 nM respectively, whereas F5 failed to bind hPD-L1 protein on Octet (Fig. 2C). To further examine whether B2 was able to functionally block the interaction between human PD-1 (hPD-1) and hPD-L1, we developed a blocking assay based on BLI technology. As shown in Fig. 2D, B2 partially blocked the interaction of hPD-1 to hPD-L1 compared with both F5 and PBS control. Moreover, B2 showed specific binding to hPD-L1 but not human B7-H3, which is another B7-CD28 family member (Fig. 2E).

**Figure 2.**
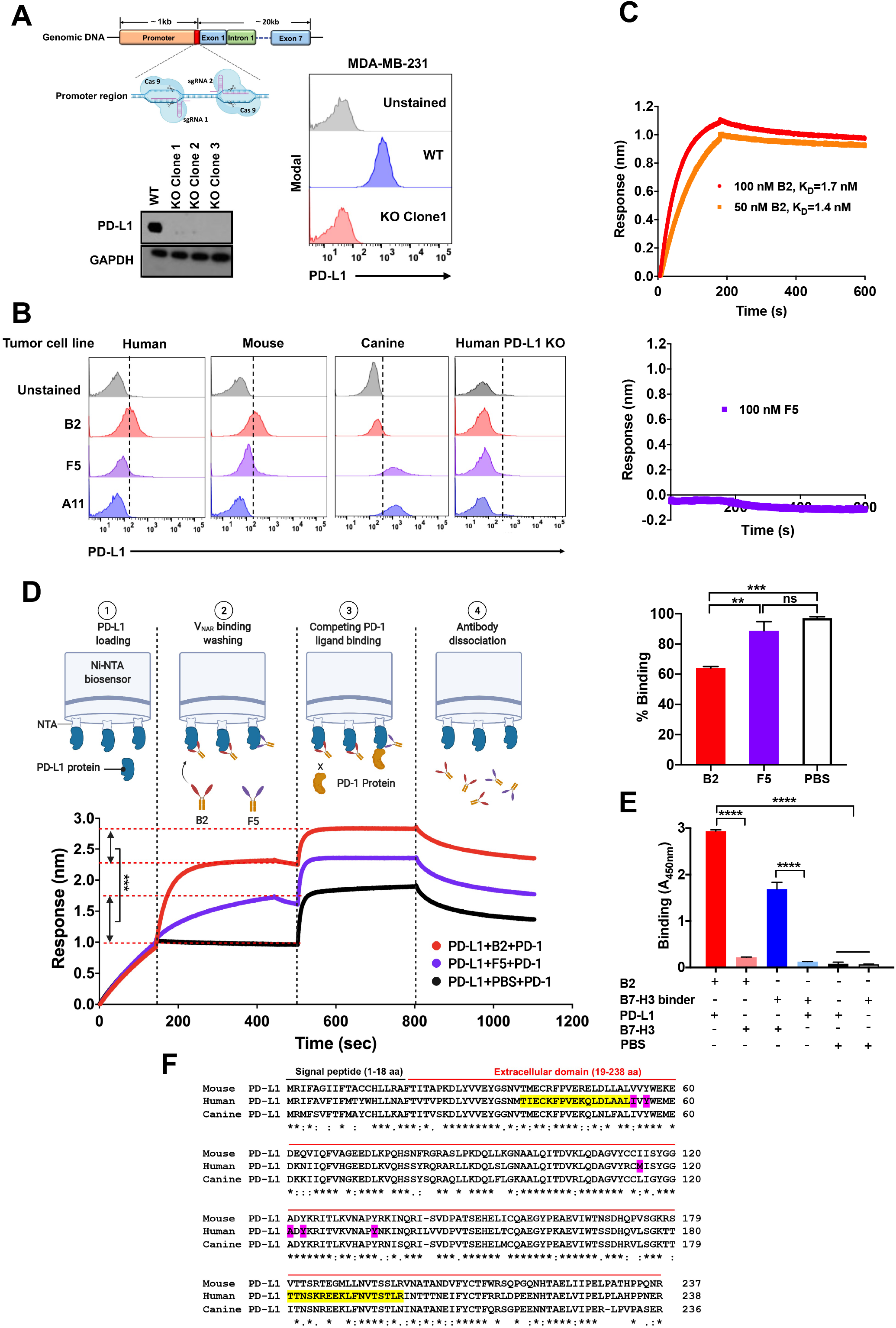
Verification of specific binding and blocking ability of anti-PD-L1 shark V_NAR_s. (A) Schematic design for constructing PD-L1 KO MDA-MB-231 cell line using CRISPR-Cas9 method. Two sgRNAs were designed to target the promoter of the endogenous PD-L1 gene. Single PD-L1 KO clones were validated by western blot and flow cytometry. (B) The cross-reactive binding of anti-PD-L1 V_NAR_s to native PD-L1 as determined by flow cytometry. Three different tumor cell lines from human, murine, and canine were stained with V_NAR_s. (C) Binding kinetics of V_NAR_-hFc to hPD-L1 protein. (D) Blocking the activity of V_NAR_-hFc to the interaction of hPD-L1 and hPD-1 as determined by the Octet platform. (E) Specific binding of B2 to hPD-L1 and hB7-H3. (F) Epitope mapping of individual B2, F5, and A11. Sequence alignment of PD-L1 ECD region of human, murine, and canine. The conserved residues are marked with asterisks (*), the residues with similar properties between variants are marked with colons (:) and the residues with marginally similar properties are marked with periods(.). The main binding residues of the hPD-L1 identified previously that interact with PD-1 are shaded in magenta. The binding peptides of B2 to hPD-L1 are highlighted in yellow. Values represent mean ± SEM. **, P < .01; ***, P < .001; ****, P < .0001; ns, not significant.

To explore the binding epitope of anti-PD-L1 nanobodies, we synthesized a peptides array based on hPD-L1 extracellular domain (ECD) that consists of total 24 peptides. As shown in Fig. S1 and 2F, both F5 and B2 significantly bind to the same peptide #19 (TTNSKREEKLFNVTSTLR), while A11 did not bind to any peptides. In comparison with F5, B2 showed specific binding to peptide #4 (TIECKFPVEKQLDLAALI), which overlaps with the PD1/PD-L1 binding site on the final amino acid “I”.

Altogether, we have successfully identified functionally cross-species anti-PD-L1 shark single-domain antibodies with high affinity.

### PD-L1 (B2) CAR-T cells kill breast cancer cells

Flow cytometric analysis showed that PD-L1 was highly expressed in multiple human tumor types, including breast cancer (MDA-MB-231), ovarian cancer (IGROV-1, OVCAR8, and NCI-ADR-RES), pancreatic cancer (KLM1 and SU8686), and lung cancer (EKVX), suggesting that PD-L1 is a putative pan-cancer antigen (Fig. 3A). To determine whether our shark V_NAR_s can be used for the CAR-T therapeutic approach, we constructed CARs containing the B2 V_NAR_ as the antigen recognition region, along with 4-1BB, CD3ζ signaling domains, and a truncated human EGFR cassette to gauge transduction efficiency and to switch CAR off (Fig. 3B). The transduction efficiency of V_NAR_ based CAR T cells was high (~90%) (Fig. 3C). During days 7–12, non-transduced mock T cells and CAR (B2) T cells showed indistinguishable expression of exhaustion markers (PD-1 and TIM-3) compared with each other, whereas slightly higher expression of LAG-3 was found in CAR (B2) T than mock T cells (Fig. 3D). MDA-MB-231 is a highly aggressive, invasive, and poorly differentiated TNBC cell line with limited treatment options. We, therefore, used it as a tumor model by engineering it to overexpress GFP/Luciferase (GL) for a luciferase-based cytolytic assay. Both mock T and CAR (B2) T cells were incubated with MDA-MB-231 GL cells for 24 hours or 96 hours. As shown in Fig. 3E, tumor cells were effectively lysed by CAR (B2) T cells in a 2-fold dose-dependent manner compared with mock T cells. Moreover, the long incubation time of 96-hours could efficiently increase the cytotoxicity of CAR (B2) T cells even at the lowest Effector: Target (E/T) ratio of 1:3. To investigate whether the cytolytic activity of CAR (B2) T cells is antigen-dependent, we incubated CAR (B2) cells with its corresponding PD-L1 KO cell line, showing that CAR T cells were not capable of killing antigen KO cells (Fig. 3E). A significantly higher level of TNF-α, IL-2, and IFN-γ was released from CAR T cells when co-cultured with tumor cells at 5:1 or 2.5:1 E/T ratios, while minimum cytokine production was observed from mock T cells (Fig. 3F). These results suggested that V_NAR_-derived T cells were able to efficiently lyse tumor cells. Furthermore, we included a corresponding soluble B2 V_NAR_ in the co-culture setup to detect whether it could affect the cytotoxicity of CAR (B2) T via blocking the recognition site on tumor cells competitively. As shown in Fig. 3G, inclusion of the B2 single domain significantly inhibited the cytolytic activity of CAR (B2) T cells. In contrast, no specific lysis in tumor cells was found in either coincubation with mock T cells or tumor cells alone in the presence of B2. Taken together, we concluded that CAR (B2) T cells could specifically lyse PD-L1 positive human tumor cells.

**Figure 3.**
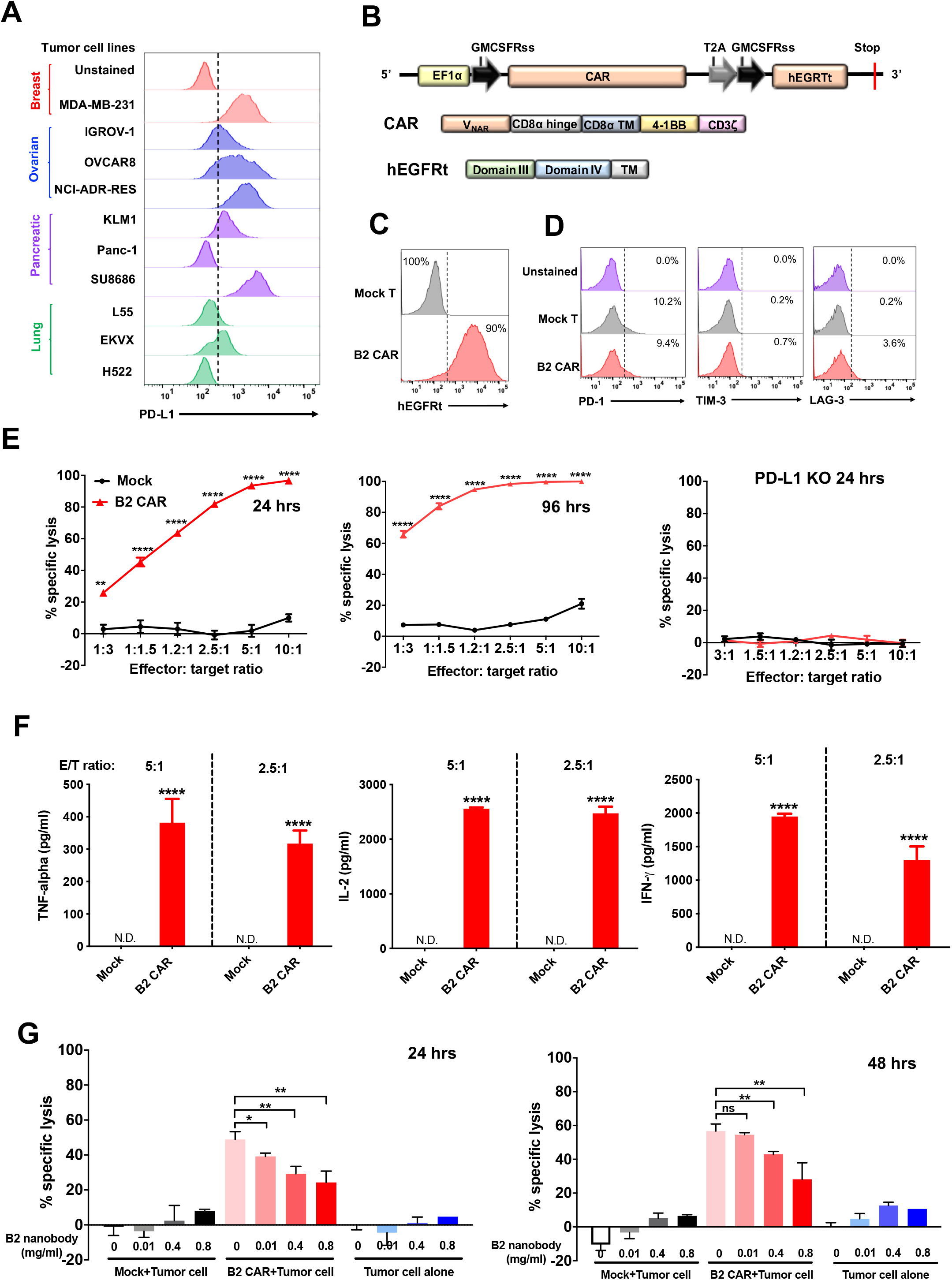
PD-L1 specific V_NAR_-based CAR-T cells exhibit antigen specific cytotoxicity against MDA-MB-231. (A) Surface PD-L1 expression on multiple human tumor types as determined by flow cytometry. (B) Construct of PD-L1 specific B2 V_NAR_-based CAR-T cell where CAR and hEGFRt are expressed separately by the self-cleaving T2A ribosomal skipping sequence. (C) The transduction efficiency of CAR (B2) in T cells was determined by hEGFRt expression. Non-transduced T cell was the mock control. (D) Exhaustion marker expression on *in vitro* cultured mock T and CAR (B2) T cell populations. (E) Cytolytic activity of CAR (B2) T cells after 24 or 96 hours of incubation with MDA-MB-231 GL or PD-L1 KO MDA-MB-231 GL respectively in a 2-fold dose dependent manner. (F) TNF-α, IL-2, and IFN-γ concentration in the supernatants of killing assay at E/T ratios of 5:1 and 2.5:1 in Fig. 3D as measured by ELISA. (G) Monovalent B2 nanobody specifically inhibited killing of CAR (B2) T cells on MDA-MB-231 cells after 24 hours and 48 hours of incubation. Tumor cells alone or mock T cells incubation in the presence of B2 nanobody were used as the control in this study. Statistical analyses are shown from three independent experiments. Values represent mean ± SEM. **, P < .01; ***, P < .001; ****, P < .0001; ns, not significant.

### CAR (B2) T cells inhibit orthotopic breast cancer in mice

To evaluate anti-tumor efficacy of CAR (B2) T cells in mice, we established an orthotopic breast tumor xenograft model via implanting the MDA-MB-231 GL line into the fourth mouse mammary fat pad. Seventeen days after tumor inoculation, mice were intravenously (IV) infused with either CAR (B2) T cells or antigen-mismatched CAR (CD19) T cells (Fig. 4A). We used both bioluminescence intensity and tumor volume to track the antitumor efficacy of CAR T cells. Mice were followed up to 8 weeks post CAR-T cell infusion except three mice from the control CAR (CD19) group or CAR (B2) treatment group that were euthanized at week 3. As shown in Fig. 4B and 4C, CAR (B2) T cells dramatically reduced breast tumor burden without a marked loss of body weight (Fig. 6D). Importantly, after 5 weeks of CAR-T infusion, we found that tumors metastasized in the control CAR (CD19) group (Fig. 4B and 4E). In contrast, no tumors metastases were found in the liver or lungs of mice that were treated with CAR (B2) T cells (Fig. 4B and 4E), indicating that CAR (B2) T cells were able to treat metastatic lesions. To determine CAR-T persistence, we recovered both CAR (CD19) and CAR (B2) T cells from mouse spleen. We found that *ex vivo* CAR (B2) T cells recovered from mice had a comparable persistence after 3 weeks infusion (Fig. 4F). Importantly, these spleen-isolated CAR (B2) T cells still showed significant *ex vivo* cytotoxicity against PD-L1 positive tumor cells compared to KO cells (Fig. 4G).

**Figure 4.**
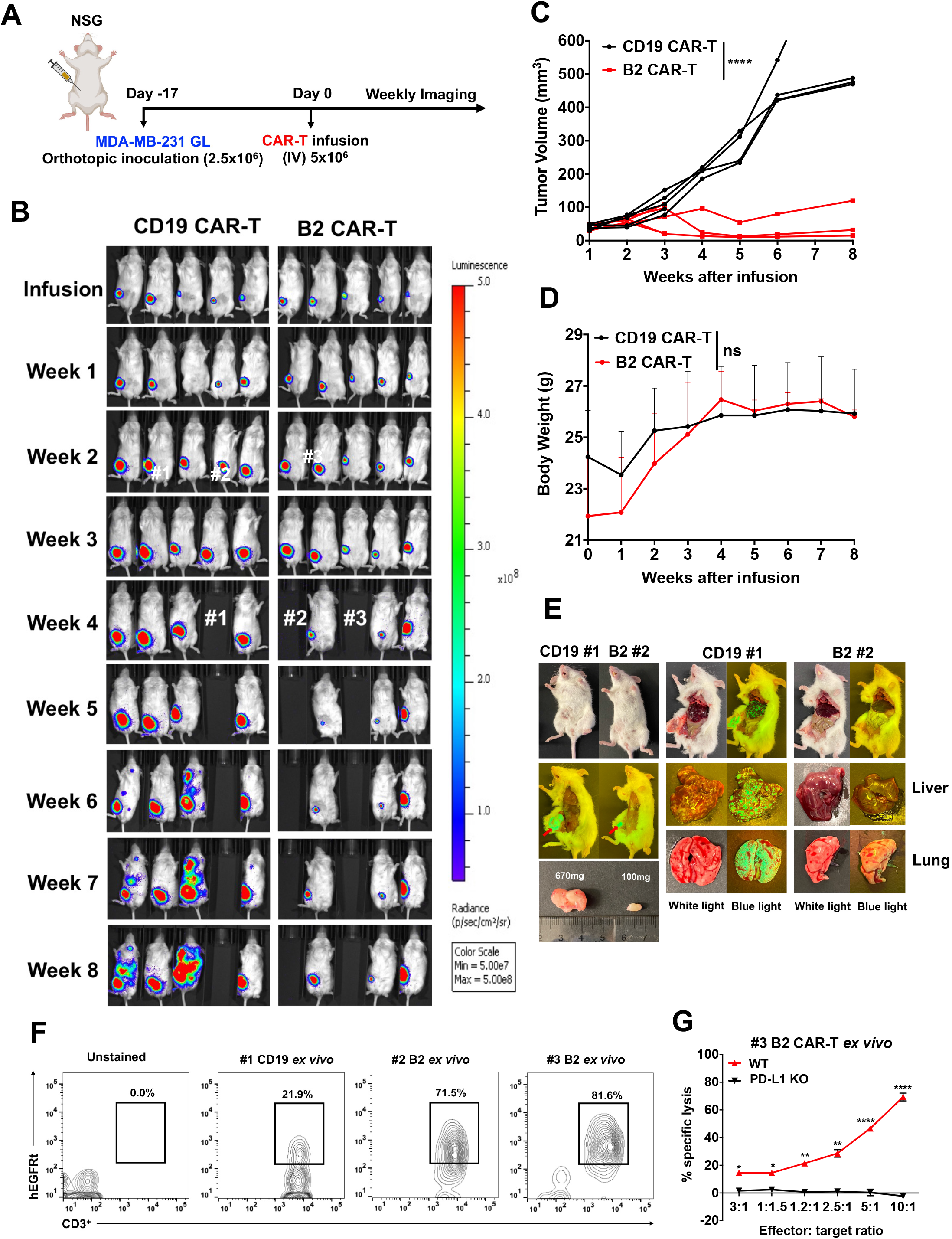
Tumor regression in the orthotopic MDA-MB-231 xenograft mouse model by CAR (B2) T cells infusion. (A) Schema of the MDA-MB-231 orthotopic xenograft NSG model IV infused with 5 million CAR (B2) T cells and CAR (CD19) CAR T cells after 17 days of tumor inoculation. (B) Representative bioluminescence image of MDA-MB-231 tumor growth in the orthotopic model. (C) Tumor size of every mouse measured by a digital caliper [V=1/2(length width^2^)]. ****, P < .0001. (D) Body weight of mice. Values shown represent mean ± SEM. (E) Representative pictures showing the restriction of tumor metastasis in CAR (B2) T cell infusion mice. (F) CAR (B2) T cell persistence and (G) *ex vivo* killing on MDA-MB-231 tumor cells after 3 weeks of CAR-T cell infusion.

### CAR (B2) T cells kill liver cancer cells by targeting inducible expression of PD-L1

Inducible but not the constitutive expression of PD-L1 can be found in liver cancer cell line Hep3B upon co-incubation with CAR (B2) T cells (Fig. 5A) possibly as a consequence of massive IFN-γ released from co-cultured CAR (B2) T (Fig. 5B). To test anti-tumor effect of CAR (B2) T to mimic the suppressive TME, we established the Hep3B xenograft mouse model with intraperitoneal (IP) injection of Hep3B GL tumor cells. After 12 days of tumor inoculation, mice were infused IP with CAR-T cells (Fig. 5C). We found that four out of five CAR (B2) T mice showed a significant decrease in tumor growth compared with the control CAR (CD19) T group after 3 weeks of infusion (Fig. 4D and E). Based on this observation, we think that CAR (B2) T cells might provide a benefit in liver cancer therapy.

**Figure 5.**
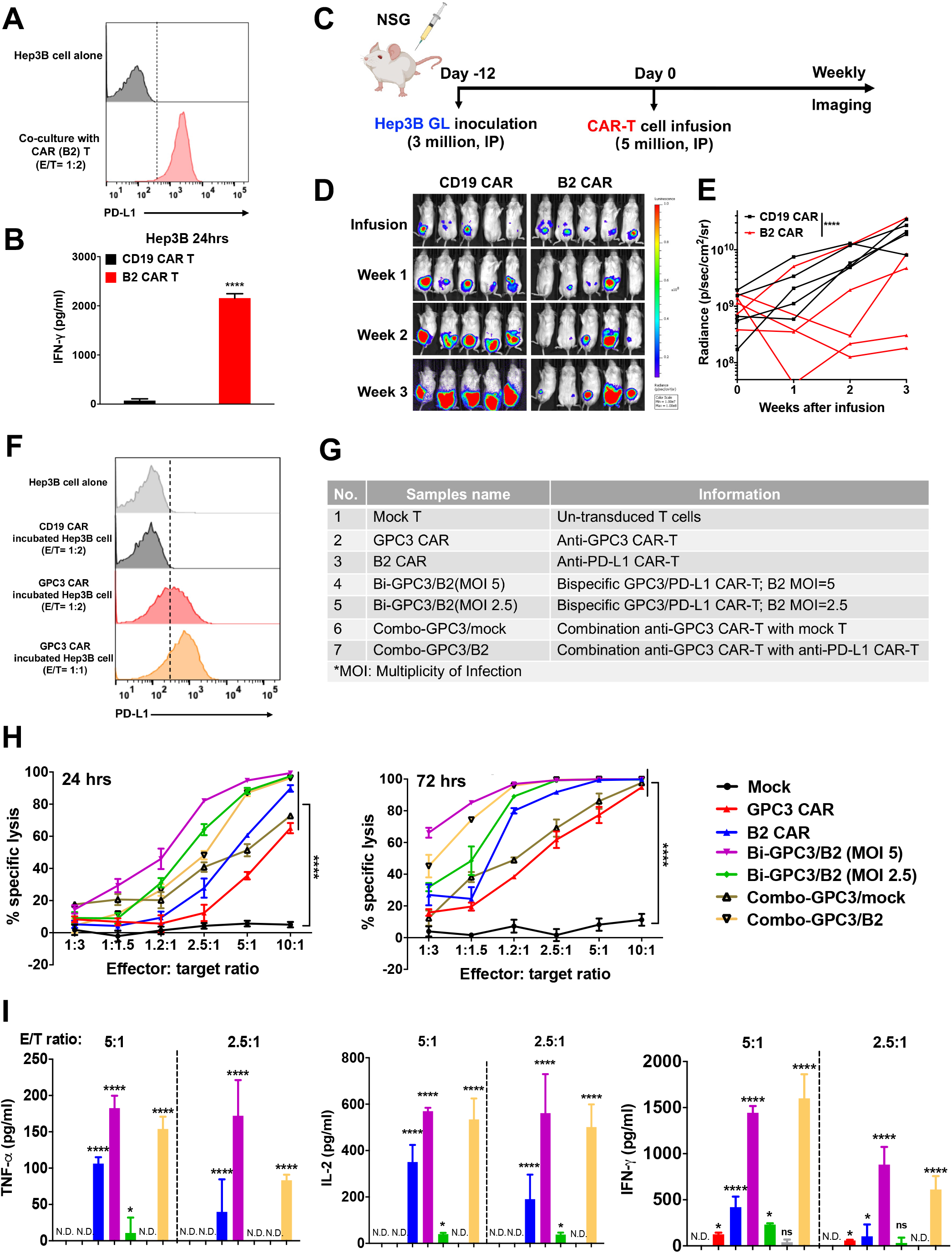
CAR (B2) T cells lysed inducible PD-L1 positive Hep3B cells and improved *in vitro* killing as engineered bispecific CAR or combination strategy with CAR-T targeting GPC3. (A) Inducible PD-L1 expression in the Hep3B tumor cells after 24 hours incubation with CAR (B2) T at E/T ratio of 1:2. (B) IFN-γ level in the supernatants of incubation CAR (CD19) T or CAR (B2) T cells with Hep3B cell. (C) Schema of the Hep3B xenograft NSG model IP infused with 5 million CAR (B2) T cells and CAR (CD19) T cells after 12 days of tumor inoculation. (D) Representative bioluminescence image of Hep3B tumor growth in the xenograft model. (E) Tumor bioluminescence growth curve. (F) Inducible PD-L1 expression in the Hep3B tumor cells alone or in the Hep3B tumor cells after 24-hours incubation with CAR (GPC3) T at E/T ratio of 1:2 or 1:1. (G) Applicable strategy of bispecific CAR-T cells and combination CAR-T cells targeting GPC3 or PD-L1. (H) Cytolytic activity of engineered CAR-T cells on Hep3B cells after 24 hours or 72 hours incubation *in vitro*. (I) TNF-α, IL-2, and IFN-γ concentration in the co-culture supernatant from (H) as measured by ELISA. Values represent mean ± SEM. **, P < .01; ***, P < .001; ****, P < .0001; ns, not significant.

### CAR (B2) T cells improve the killing effect of CAR (GPC3) T cells in liver cancer

In our previous study, we developed GPC3-targeted CAR-T cells as an emerging liver cancer therapy^10^. We observed that CAR (GPC3) T cells killed Hep3B tumor cells efficiently but upregulated PD-L1 expression was found in CAR (GPC3) T-cocultured Hep3B cells (Fig. 5F), which may allow cancers to evade the host immune system. Therefore, we hypothesized that the elimination of inducible PD-L1 positive tumor cells by CAR (B2) T cells will improve the anti-tumor activity. To detect our hypothesis, we designed two strategies, including bispecific expression or combination of PD-L1 and GPC3 CAR-T cells (Fig. 5G). Bispecific CAR-T cell was produced by co-transducing with GPC3 CAR and CAR (B2) lentivirus (Fig. 5G). To compare their anti-tumor effect, all seven groups of CAR-T cells and mock T cells (Fig. 5G) were incubated with Hep3B cells for 24 hours and 72 hours. As shown in Fig. 5H, the cytotoxicity of bispecific CARs was significantly higher than either of the monospecific CARs, especially at 72 hours of incubation time. Moreover, CAR (B2) T cells can improve the efficiency of CAR (GPC3) T cells in a dose-dependent manner (MOI 2.5 vs 5). Furthermore, we observed higher levels of TNF-α, IL-2, and IFN-γ were secreted from both bispecific and combination CAR treatment than those of monospecific CAR-T treatments (Fig. 5I). Therefore, we concluded that the bispecific CAR-T and the combined CAR-T strategies significantly improved the activity of CAR-T cell in liver cancer by targeting both PD-L1 and GPC3.

### Combination of CAR (B2) T and CAR (GPC3) T achieves a synergistic anti-tumor effect in mice

To further analyze the functions of bi-specific CAR-T and combination CAR-T strategies in response to liver cancer, we confirmed the anti-tumor effect through the Hep3B xenograft mouse model. Mice bearing Hep3B tumors were divided into five groups and infused with 5 million equivalents of CAR (GPC3) T, CAR (CD19) T, CAR (B2) T, Bi-GPC3/B2 CAR T, and a combination of 2.5 million CAR (GPC3) T and 2.5 million CAR (B2) T (referred to as “Combo-GPC3/B2”) cells, respectively. Tumor luciferase signal was evaluated by bioluminescence imaging weekly, and T cells isolated from week 2 mouse blood were analyzed (Fig. 6A). In comparison with the control CAR (CD19) T cells, CAR (GPC3) T and CAR (B2) T cells individually inhibited tumor growth in xenografts (Fig. 6B and 6C). Surprisingly, bispecific CAR-T cells failed to regress tumor burden and the effect was worse compared to monospecific CAR-T cells. However, the combination group showed a significant synergistic anti-tumor effect in xenografts (Fig. 6B and 6C). We sacrificed mice by the end of week 4 after treatment due to maximum tumor limitation. To visualize tumor size, we isolated tumors from a mouse (#1) from combination group and a mouse (#2) from bispecific CAR-T group. As shown in Fig. 6D, the tumor size from combination group was much smaller than that from bispecific CAR treatment group. To identify factors that contribute to the high efficiency in combination CAR-T strategy, we detected number, immunophenotype, and exhaustion of CAR-T cells isolated from mouse blood at week 2 of infusion. We found that mice receiving CAR (B2) T, Combo-GPC3/B2 CAR T, or Bi-GPC3/B2 CAR T cells had much higher CD3+CAR+ T cells counts in blood than those who received CAR (CD19) T or CAR (GPC3) T cells (Fig. 6E). On the other hand, the number of CAR (B2) T was higher than that of combination followed by bispecific CAR-T cells, indicating bispecific CAR-T might loss PD-L1-specific proliferation. Indeed, we found that the recovered CAR-T cells from Combo-GPC3/B2 mouse (#143) showed a higher binding ability to PD-L1 than the CAR-T cells of Bi-GPC3/B2 mouse (#107) (21.2% vs 16.4%) though both CAR-T groups showed similar binding percentage in cell culture (3.28% vs 2.62%) (Fig. 6F). Moreover, the CAR-T cells recovered from the mouse spleen showed higher binding ability compared with *in vitro* cultured CAR-T cells, especially on CAR (B2) T cells (5.93% vs 35.4%). Besides the functional capacity of endogenous T cells, the frequency of memory T cell subset is also associated with tumor response. Here, we analyzed the T differentiation subsets consisting of stem cell-like memory T cells (T_SCM_: CD62L+CD45RA+CD95+), central memory T cells (T_CM_: CD62L+CD45RA-CD95+), effector memory T cells (T_EM_: CD62L-CD45RA-CD95+), and terminally differentiated effector memory T cells (T_EMRA_: CD62L-CD45RA+CD95+) in CD4+CAR+ and CD8+CAR+ subpopulations in mouse blood after 2 weeks of infusion. As shown in Fig. 6G, the combination group exhibited a significantly higher percentage of Tscm than CAR (B2) T and higher frequency of Tem and Tcm than CAR (GPC3) T on both CD4+ and CD8+ subpopulations. We further analyzed the expression of co-inhibitory receptors in CAR-T cells, including PD-1, LAG-3, and TIM-3. The CAR-T cells that containing B2 showed higher expression of PD-1 and LAG-3 than CAR (GPC3) in both CD4+ and CD8+ subpopulations (Fig. 6H). Collectively, these results suggest that a combination of CAR (GPC3) T and CAR (B2) T, but not bispecific CAR, synergistically killed Hep3B tumor.

**Figure 6.**
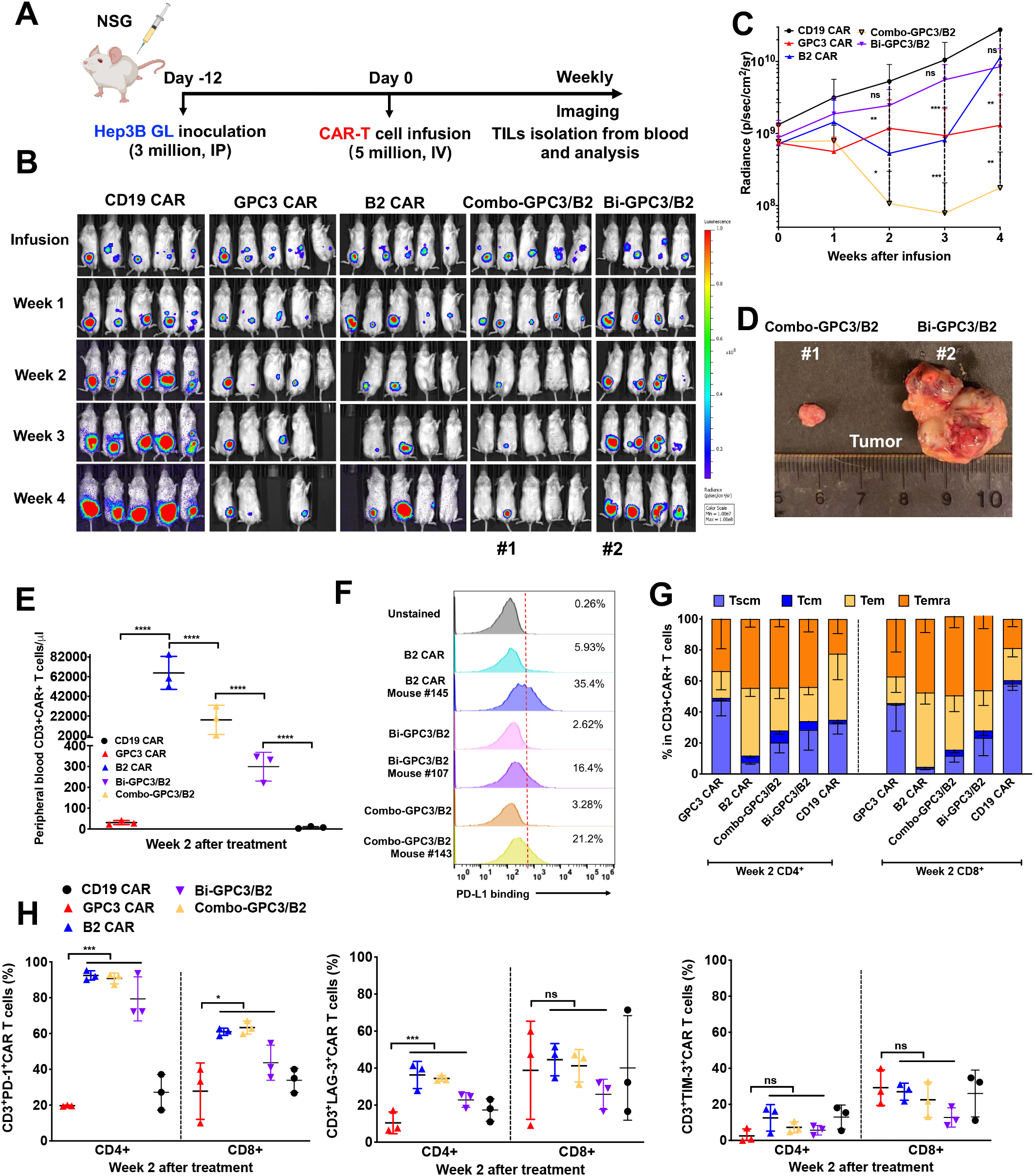
Combined CAR (B2) T with CAR (GPC3) T cells achieve a synergistic anti-tumor effect *in vivo*. (A) Schema of the Hep3B xenograft NSG model IP infused with equivalent 5 million CAR T cells after 12 days of tumor inoculation. (B) Representative bioluminescence image of Hep3B tumor growth in the xenograft model (C) Tumor bioluminescence growth curve. (D) The sizes of tumors in mice from combination CAR group (#1 mouse) and bispecific group (#2 mouse) at the end of the study. (E) Absolute CAR-T count was detected in mouse peripheral blood after 2 weeks of treatment. Absolute CAR-T concentration (cells/μL) ±SD for all evaluable mice in each treatment group is shown. (F) The binding ability of both *in vitro* and *in vivo* recovered CAR-T cells to PD-L1 antigen using flow cytometry. (G) Relative proportion of stem cell-like memory (T_SCM_), central memory (T_CM_), effector memory (T_EM_), and terminally differentiated effector memory (T_EMRA_) subsets defined by CD62L, CD45RA and CD95 expression in both CD4 + and CD8+ CAR+T cell population in mouse blood on week 2 of treatment. (H) Exhaustion marker expression on CD4 + and CD8 + CAR+T cells populations in mouse blood on week 2 of treatment.

## Discussion

Checkpoint molecule PD-L1 is highly expressed on many tumors in a constitutive or IFN-γ-inducible manner. IFN-γ is the key functional cytokine released from effector T cells; however, the increased expression of PD-L1 on tumor cells binding to PD-1 on effector T cells results in T cell exhaustion, and inhibition of T cell functions^30^. In this study, we hypothesized that the development of CAR-T cells targeting PD-L1 could kill solid tumors via recognizing the constitutive or inducible expression of PD-L1 in the tumor immunosuppressive microenvironment. To test our hypothesis, we isolated a panel of anti-PD-L1 single domain antibodies from a newly established semi-synthetic nurse shark V_NAR_ library. The best candidate, B2, showed a specific binding ability to PD-L1, and was cross-reacting with both human and mouse antigens. Importantly, B2 functionally blocked the interaction between PD-L1 and PD-1. Moreover, we found that single domain-based CAR-T showed much higher transduction efficiency than scFv-based CAR-T cells, indicating that single domain antibodies are more appropriate to be engineered into CAR format because it is smaller, easily expressible, and more stable.

PD-L1 is not only overexpressed on a larger number of malignancies, but also on immune cells in the tumor microenvironment^13^. T cells express low levels of endogenous PD-L1, which leads to the development of CAR-T cells that targeting PD-L1 is somewhat intricate by killing PD-L1 expressing tumor cells and blocking the PD-1/PD-L1 checkpoint axis^31,32^. Antigen exposure of CAR-T cells may lead to T cell fratricide and exhaustion, impairing the proliferation and persistence of CAR-T cells *in vitro* and *in vivo*. Xie *et al*. reported that camelid V_H_H-based anti-mouse PD-L1 CAR-T cells can “self-activate *in vitro* and PD-L1 deficient CAR-T cells could live longer than WT CAR-T^31^. However, during a period of 7-12 days *in vitro* co-culturing with CD3/CD28 microbeads, we did not find upregulation in PD-L1 (Fig. S2) or exhaustion markers (PD-1, TIM-3, and LAG-3) in the activated CAR (B2) T cells compared with mock T cells *in vitro*. These events were probably due to PD-L1 antigen endocytosis caused by anti-PD-L1 CAR-T cells themselves^33,34^. Interestingly, we did not observe any cytolytic phenomenon or upregulated IFN-γ expression in the cultured CAR (B2) T cells, indicating that the cytotoxicity of CAR (B2) T was not triggered by the T cells’ endogenous PD-L1. We consider that less tonic signaling of our shark V_NAR_-based CAR (B2) T cells may be due to the relative low binding affinity of CAR (B2). Ghorashian *et al*. reported the enhanced proliferation and anti-tumor activity in a lower affinity CD19 CAR comparing with that in a clinical high affinity CD19 CAR-T, indicating that the increased immunoreceptor affinity may adversely affect T cell reponses^35^.

To overcome tumor escape mechanisms and enhance the anti-tumor effect of CAR-T cells, a combination strategy might be more feasible in solid tumor therapy, such as combining CAR-T cells with monoclonal antibodies, small-molecules, or bi-specific CAR T cells targeting different tumor-specific antigens^36,37^. In our study, we found that CAR (B2) T cells could kill liver cancer cells by targeting inducible PD-L1 in the immunosuppressive TME (Fig. 5D), whereas B2 V_NAR_ did not show a significant benefit in improving cytotoxicity of CAR (GPC3) T cell even though it functionally blocked the interaction of PD-1 to PD-L1 (Fig. S3). Thus, we constructed the bispecific CAR-T cell targeting both HCC tumor-specific antigen GPC3 and inducible tumor-immunosuppressive antigen PD-L1. Surprisingly, Bi-GPC3/B2 CAR T cells worked best *in vitro* whereas only slightly inhibited liver tumor progression *in vivo*, and even worse than the individual CAR (GPC3) T and CAR (B2) T cells. We think it may be due to low CAR density or low binding affinity when we co-transduced B2 and GPC3 CAR lentivirus into PBMCs. In future work, we may optimize the bi-specific CAR construct by optimizing GPC3 and B2 CAR fragments into one construct^19^. Encouragingly, the combination of CAR (GPC3) T and CAR (B2) T cell achieved a synergistic anti-tumor effect *in vivo*. A previous study reported that the combination of anti-mesothelin CAR-T cell with anti-PD-L1 CAR-T did not repress tumor growth synergistically in PDX, as anti-PD-L1 CAR-T killed anti-mesothelin CAR-T cell by targeting its endogenous PD-L1 antigen^33^. In our study, we did not observe upregulated PD-L1 expression in CAR (GPC3) T cells probably due to different CAR constructs. On the other hand, we found that the expansion of CAR-T count in mouse blood is highly correlated with the presence of CAR (B2) construct. It may be due to the cross-recognition of CAR (B2) to mouse antigen, but the CAR-T treatment mice were healthy and did not experience body weight loss, indicating our CAR (B2) T cells are safe for the mouse. Although we didn’t observe upregulated PD-1 expression in the cultured CAR (B2) T cells, the high expression of PD-1 was found in *ex vivo* B2-related CAR-T cells (Fig. 6G). However, the recovered CAR (B2) T cells from 3 weeks after infusion still efficiently lysed the MDA-MB-231 cells (Fig. 6G), probably due to the B2 V_NAR_ blocking the interaction of PD-1 to PD-L1 even though not entirely.

## Conclusions

We have demonstrated that a semi-synthetic shark V_NAR_ phage library based on fully randomized CDR3 can be used in isolating anti-PD-L1 specific single domain antibodies. We conclude that the PD-L1-targeted shark V_NAR_-based CAR-T cell is a promising strategy in triple-negative breast cancer and liver cancer therapy, providing a rationale for the potential use of PD-L1 (B2) CAR-T cells in clinical studies. Overall, the results in this study demonstrate the feasibility and the efficacy of CAR-T cells targeting tumor immunosuppressive microenvironment antigen PD-L1 against aggressive solid tumors. To improve treatment of solid tumors, future efforts should be directed at utilizing genome editing to develop “off-the-shelf” fratricide-resistant PD-L1-targeted CAR-T cells lacking both endogenous PD-L1 and T cell receptor alpha chain expression on T cells.

## Materials and Methods

### Construction of a synthetic 18AA CDR3 nurse shark V_NAR_ phage library

We constructed the new synthetic 18AA CDR3 nurse shark V_NAR_ phage library based on our previous naïve shark library^26^. For the V_NAR_s DNA cassettes, a non-canonical cysteine in CDR1 was mutated to tyrosine (C29Y) using naïve shark library V_NAR_s pComb3x plasmid as the template. Subsequently, a pair of randomized 18AA CDR3 primers was designed to amplify the CDR3 loop using the PCR method. PCR product were circularized by intra-molecular self-ligation in 1 ml of ligation buffer using T4 DNA ligase (New England Biolabs, Ipswich, MA). Finally, the ligation products were purified by removing the enzymes and transformed into 500 μl of electroporation competent TG1 cells (Lucigen, Middleton, WI) to make the library.

### Phage panning

The phage panning protocol has been described previously^26,38^. The mPD-L1 protein bought from R&D Systems was used for four rounds of panning. Details are provided in the supplemental materials.

### Affinity binding and blocking activity

The binding kinetics of the V_NAR_-hFc (produced by GenScript) to hPD-L1-His protein (SinoBiological) was determined using the Octet RED96 system (FortéBio) at the Biophysics Core (National Heart, Lung and Blood Institute or NHLBI) as described previously^39^. The blocking activity of B2-hFc was also determined using the BLI Octet platform as described previously^40^. Details are provided in the supplemental materials.

### Generation of anti-PD-L1 nanobody-based CAR-T cells

We generated the PD-L1-target shark V_NAR_-based CAR-T lentiviral vector following the design principle of CAR construct published in our previous study^10^. Briefly, the V_NAR_ fragment of B2 was subcloned into a CAR construct (pMH330). The CAR expressing lentivirus was produced as described previously^10^. PBMCs isolated from healthy donors were stimulated for 24h using anti-CD3/anti-CD28 antibody-coated beads (Invitrogen) at a bead: cell ratio of 2:1 according to manufacturer’s instructions in the presence of IL-2.

### *In vitro* cytolysis of CAR-T cells and activation assays

The cytotoxicity of CAR-T cells was determined by a luciferase-based assay. In brief, the luciferase-expressing MDA-MB-231 and Hep3B tumor cells were used to establish a cytolytic assay. The cytolysis of PD-L1-target CAR (B2) T cells was detected by co-culturing with MDA-MB-231 GFP-Luc and Hep3B GFP-Luc at various E/T ratios for 24 hours or 96 hours followed by measurement of the luciferase activity using the luciferase assay system (Promega) on Victor (PerkinElmer). The supernatants were collected for TNF-α, IL-2, and IFN-γ detection using ELISA Kit (BD biosciences). In the killing blocking assay of CAR-T cells, varying concentration of soluble B2 protein was added into tumor CAR-T cells incubation for 24 hours and 48 hours.

### Animal studies

5-week-old female NOD/SCID/IL-2Rgc^null^ (NSG) mice (NCI Frederick) were housed and treated under the protocol (LMB-059) approved by the Institutional Animal Care and Use Committee at the NIH. A total of 3 million MDA-MB-231-GFP-Luc cells were suspended in the mixture of PBS: Matrigel (BD Biosciences) at 1:1, and inoculated into the inguinal mammary fat pad to establish the orthotopic MDA-MB-231 model. Peritoneal Hep3B xenograft tumor model was established as previously described^10^. Tumor volume was calculated as ½ (length × width^2^) and bioluminescent intensity (Xenogen IVIS Lumina). When the average tumor size reached the indicated size, 5 million CAR-T cells were intravenously injected into mice models. *Ex vivo* T cells were isolated from mice spleens using Miltenyi Biotec tumor dissociation kit, and were cultured *in vitro* with 40ng/ul IL-2, IL-7, and IL-21 in the culture media.

### Statistical analysis

All experiments were repeated at least three times to ensure reproducibility of results. All statistical analyses were performed using GraphPad Prism, and are presented as mean±SEM. Results were analyzed using 2-tailed unpaired Student’s t test. A P value of < 0.05 was considered statistically significant.

## Acknowledgments

We thank NIH Fellows Editorial Board for editorial assistance. We thank NCI CCR Animal Resource Program/NCI Biological Testing Branch, NCI CCR/Leidos Animal Facility, Drs. Grzegorz Piszczek and Di Wu of the NHLBI Biophysics Core, NCI CCR Flow Cytometry Core Facility for providing assistance in animal support, V_NAR_ single domain antibody kinetics/affinity analysis and cellular staining.

## Authors’ Contributions

Conception and design, D.L., G.M., and M.H.; development of methodology, D.L., H.J.E., G.M., C.P.D., and M.H.; acquisition of data, D.L., H.J.E., J.H., T.Y.Z.L.; analysis and interpretation of data, D.L., H.J.E., J.H., T.Y.Z.L.; writing, D.L. and M.H.; review and/or revision of manuscript, T.Y.Z.L., G.M., C.P.D., and M.H.; study supervision, M.H. All authors read and approved the final manuscript.

## Funding

This work was supported by the NCI CCR FLEX Program Synergy Award (to G.M. and M.H.) and the NCI CCR FLEX Program Technology Development Award (to M.H.).

## Competing interests

M.H., G.M., D.L., H.J.E., and C.P.D. are inventors on US provisional patent application no. 63/208,755, “Cross Species Single Domain Antibodies Targeting PD-L1 For Treating Solid Tumors”. The authors declare no other competing interests.

## Ethics approval

All mice were housed and treated under the protocol (LMB-059) approved by the Institutional Animal Care and Use Committee at the NIH.

## Provenance and peer review

Not commissioned, externally peer reviewed.

## Data availability statement

All data relevant to the study are included in the article or uploaded as online supplemental information.

## References

1. Rosenberg, S.A., Restifo, N.P., Yang, J.C., Morgan, R.A., and Dudley, M.E. (2008). Adoptive cell transfer: a clinical path to effective cancer immunotherapy. Nat Rev Cancer 8, 299–308. 10.1038/nrc2355.

2. Rosenberg, S.A., and Restifo, N.P. (2015). Adoptive cell transfer as personalized immunotherapy for human cancer. Science 348, 62–68. 10.1126/science.aaa4967.

3. Kochenderfer, J.N., Wilson, W.H., Janik, J.E., Dudley, M.E., Stetler-Stevenson, M., Feldman, S.A., Maric, I., Raffeld, M., Nathan, D.A., Lanier, B.J., Morgan, R.A., et al. (2010). Eradication of B-lineage cells and regression of lymphoma in a patient treated with autologous T cells genetically engineered to recognize CD19. Blood 116, 4099–4102. 10.1182/blood-2010-04-281931.

4. Maher, J., Brentjens, R.J., Gunset, G., Riviere, I., and Sadelain, M. (2002). Human T-lymphocyte cytotoxicity and proliferation directed by a single chimeric TCRzeta /CD28 receptor. Nat Biotechnol 20, 70–75. 10.1038/nbt0102-70.

5. Imai, C., Mihara, K., Andreansky, M., Nicholson, I.C., Pui, C.H., Geiger, T.L., and Campana, D. (2004). Chimeric receptors with 4-1BB signaling capacity provoke potent cytotoxicity against acute lymphoblastic leukemia. Leukemia 18, 676–684. 10.1038/sj.leu.2403302.

6. Song, D.G., Ye, Q., Carpenito, C., Poussin, M., Wang, L.P., Ji, C., Figini, M., June, C.H., Coukos, G., and Powell, D.J., Jr. (2011). In vivo persistence, tumor localization, and antitumor activity of CAR-engineered T cells is enhanced by costimulatory signaling through CD137 (4-1BB). Cancer Res 71, 4617–4627. 10.1158/0008-5472.CAN-11-0422.

7. Gross, G., Waks, T., and Eshhar, Z. (1989). Expression of immunoglobulin-T-cell receptor chimeric molecules as functional receptors with antibody-type specificity. Proc Natl Acad Sci U S A 86, 10024–10028. 10.1073/pnas.86.24.10024.

8. Kochenderfer, J.N., and Rosenberg, S.A. (2013). Treating B-cell cancer with T cells expressing anti-CD19 chimeric antigen receptors. Nature reviews. Clinical oncology 10, 267–276. 10.1038/nrclinonc.2013.46.

9. Li, N., Fu, H., Hewitt, S.M., Dimitrov, D.S., and Ho, M. (2017). Therapeutically targeting glypican-2 via single-domain antibody-based chimeric antigen receptors and immunotoxins in neuroblastoma. Proc Natl Acad Sci U S A 114, E6623–E6631. 10.1073/pnas.1706055114.

10. Li, D., Li, N., Zhang, Y.F., Fu, H., Feng, M., Schneider, D., Su, L., Wu, X., Zhou, J., Mackay, S., Kramer, J., et al. (2020). Persistent Polyfunctional Chimeric Antigen Receptor T Cells That Target Glypican 3 Eliminate Orthotopic Hepatocellular Carcinomas in Mice. Gastroenterology 158, 2250–2265 e2220. 10.1053/j.gastro.2020.02.011.

11. Lv, J., Zhao, R.C., Wu, D., Zheng, D.W., Wu, Z.P., Shi, J.X., Wei, X.R., Wu, Q.T., Long, Y.G., Lin, S.M., Wang, S.N., et al. (2019). Mesothelin is a target of chimeric antigen receptor T cells for treating gastric cancer. Journal of hematology & oncology 12. ARTN 18 10.1186/s13045-019-0704-y.

12. Zhang, Z., Jiang, D., Yang, H., He, Z., Liu, X., Qin, W., Li, L., Wang, C., Li, Y., Li, H., Xu, H., et al. (2019). Modified CAR T cells targeting membrane-proximal epitope of mesothelin enhances the antitumor function against large solid tumor. Cell death & disease 10, 476. 10.1038/s41419-019-1711-1.

13. Sun, C., Mezzadra, R., and Schumacher, T.N. (2018). Regulation and Function of the PD-L1 Checkpoint. Immunity 48, 434–452. 10.1016/j.immuni.2018.03.014.

14. Dong, H., Strome, S.E., Salomao, D.R., Tamura, H., Hirano, F., Flies, D.B., Roche, P.C., Lu, J., Zhu, G., Tamada, K., Lennon, V.A., et al. (2002). Tumor-associated B7-H1 promotes T-cell apoptosis: a potential mechanism of immune evasion. Nat Med 8, 793–800. 10.1038/nm730.

15. Weinstock, M., and McDermott, D. (2015). Targeting PD-1/PD-L1 in the treatment of metastatic renal cell carcinoma. Ther Adv Urol 7, 365–377. 10.1177/1756287215597647.

16. Sznol, M. (2014). Blockade of the B7-H1/PD-1 pathway as a basis for combination anticancer therapy. Cancer J 20, 290–295. 10.1097/PPO.0000000000000056.

17. Xie, Y.J., Dougan, M., Jailkhani, N., Ingram, J., Fang, T., Kummer, L., Momin, N., Pishesha, N., Rickelt, S., Hynes, R.O., and Ploegh, H. (2019). Nanobody-based CAR T cells that target the tumor microenvironment inhibit the growth of solid tumors in immunocompetent mice. P Natl Acad Sci USA 116, 7624–7631. 10.1073/pnas.1817147116.

18. Fabian, K.P., Padget, M.R., Donahue, R.N., Solocinski, K., Robbins, Y., Allen, C.T., Lee, J.H., Rabizadeh, S., Soon-Shiong, P., Schlom, J., and Hodge, J.W. (2020). PD-L1 targeting high-affinity NK (t-haNK) cells induce direct antitumor effects and target suppressive MDSC populations. J Immunother Cancer 8. 10.1136/jitc-2019-000450.

19. Zhao, W., Jia, L., Zhang, M., Huang, X., Qian, P., Tang, Q., Zhu, J., and Feng, Z. (2019). The killing effect of novel bi-specific Trop2/PD-L1 CAR-T cell targeted gastric cancer. Am J Cancer Res 9, 1846–1856.

20. Chailyan, A., Marcatili, P., and Tramontano, A. (2011). The association of heavy and light chain variable domains in antibodies: implications for antigen specificity. FEBS J 278, 2858–2866. 10.1111/j.1742-4658.2011.08207.x.

21. Hamers-Casterman, C., Atarhouch, T., Muyldermans, S., Robinson, G., Hamers, C., Songa, E.B., Bendahman, N., and Hamers, R. (1993). Naturally occurring antibodies devoid of light chains. Nature 363, 446–448. 10.1038/363446a0.

22. Flajnik, M.F., and Kasahara, M. (2010). Origin and evolution of the adaptive immune system: genetic events and selective pressures. Nat Rev Genet 11, 47–59. 10.1038/nrg2703.

23. Muyldermans, S. (2013). Nanobodies: natural single-domain antibodies. Annu Rev Biochem 82, 775–797. 10.1146/annurev-biochem-063011-092449.

24. Criscitiello, M.F., Saltis, M., and Flajnik, M.F. (2006). An evolutionarily mobile antigen receptor variable region gene: doubly rearranging NAR-TcR genes in sharks. Proc Natl Acad Sci U S A 103, 5036–5041. 10.1073/pnas.0507074103.

25. English, H., Hong, J., and Ho, M. (2020). Ancient species offers contemporary therapeutics: an update on shark VNAR single domain antibody sequences, phage libraries and potential clinical applications. Antibody therapeutics 3, 1–9. 10.1093/abt/tbaa001.

26. Feng, M., Bian, H., Wu, X., Fu, T., Fu, Y., Hong, J., Fleming, B.D., Flajnik, M.F., and Ho, M. (2019). Construction and next-generation sequencing analysis of a large phage-displayed VNAR single-domain antibody library from six naive nurse sharks. Antibody therapeutics 2, 1–11. 10.1093/abt/tby011.

27. Matz, H., and Dooley, H. (2019). Shark IgNAR-derived binding domains as potential diagnostic and therapeutic agents. Dev Comp Immunol 90, 100–107. 10.1016/j.dci.2018.09.007.

28. Ubah, O.C., Buschhaus, M.J., Ferguson, L., Kovaleva, M., Steven, J., Porter, A.J., and Barelle, C.J. (2018). Next-generation flexible formats of VNAR domains expand the drug platform’s utility and developability. Biochem Soc Trans 46, 1559–1565. 10.1042/BST20180177.

29. Camacho-Villegas, T., Mata-Gonzalez, T., Paniagua-Solis, J., Sanchez, E., and Licea, A. (2013). Human TNF cytokine neutralization with a vNAR from Heterodontus francisci shark: a potential therapeutic use. MAbs 5, 80–85. 10.4161/mabs.22593.

30. Chen, L.P., and Han, X. (2015). Anti-PD-1/PD-L1 therapy of human cancer: past, present, and future. Journal of Clinical Investigation 125, 3384–3391. 10.1172/Jci80011.

31. Xie, Y.J., Dougan, M., Jailkhani, N., Ingram, J., Fang, T., Kummer, L., Momin, N., Pishesha, N., Rickelt, S., Hynes, R.O., and Ploegh, H. (2019). Nanobody-based CAR T cells that target the tumor microenvironment inhibit the growth of solid tumors in immunocompetent mice (vol 116, pg 7624, 2019). P Natl Acad Sci USA 116, 16656–16656. 10.1073/pnas.1912487116.

32. Qin, L., Zhao, R.C., Chen, D.M., Wei, X.R., Wu, Q.T., Long, Y.G., Jiang, Z.W., Li, Y.Q., Wu, H.P., Zhang, X.C., Wu, Y.L., et al. (2020). Chimeric antigen receptor T cells targeting PD-L1 suppress tumor growth. Biomark Res 8. ARTN 19 10.1186/s40364-020-00198-0.

33. Qin, L., Zhao, R., Chen, D., Wei, X., Wu, Q., Long, Y., Jiang, Z., Li, Y., Wu, H., Zhang, X., Wu, Y., et al. (2020). Chimeric antigen receptor T cells targeting PD-L1 suppress tumor growth. Biomark Res 8, 19. 10.1186/s40364-020-00198-0.

34. Hamieh, M., Dobrin, A., Cabriolu, A., van der Stegen, S.J.C., Giavridis, T., Mansilla-Soto, J., Eyquem, J., Zhao, Z., Whitlock, B.M., Miele, M.M., Li, Z., et al. (2019). CAR T cell trogocytosis and cooperative killing regulate tumour antigen escape. Nature 568, 112–116. 10.1038/s41586-019-1054-1.

35. Ghorashian, S., Kramer, A.M., Onuoha, S., Wright, G., Bartram, J., Richardson, R., Albon, S.J., Casanovas-Company, J., Castro, F., Popova, B., Villanueva, K., et al. (2019). Enhanced CAR T cell expansion and prolonged persistence in pediatric patients with ALL treated with a low-affinity CD19 CAR. Nat Med 25, 1408–1414. 10.1038/s41591-019-0549-5.

36. Pan, Z., Di, S., Shi, B., Jiang, H., Shi, Z., Liu, Y., Wang, Y., Luo, H., Yu, M., Wu, X., and Li, Z. (2018). Increased antitumor activities of glypican-3-specific chimeric antigen receptor-modified T cells by coexpression of a soluble PD1-CH3 fusion protein. Cancer Immunol Immunother 67, 1621–1634. 10.1007/s00262-018-2221-1.

37. Hegde, M., Mukherjee, M., Grada, Z., Pignata, A., Landi, D., Navai, S.A., Wakefield, A., Fousek, K., Bielamowicz, K., Chow, K.K., Brawley, V.S., et al. (2016). Tandem CAR T cells targeting HER2 and IL13Ralpha2 mitigate tumor antigen escape. J Clin Invest 126, 3036–3052. 10.1172/JCI83416.

38. Ho, M., Kreitman, R.J., Onda, M., and Pastan, I. (2005). In vitro antibody evolution targeting germline hot spots to increase activity of an anti-CD22 immunotoxin. J Biol Chem 280, 607–617. 10.1074/jbc.M409783200.

39. Maus, M.V., and June, C.H. (2016). Making Better Chimeric Antigen Receptors for Adoptive T-cell Therapy. Clin Cancer Res 22, 1875–1884. 10.1158/1078-0432.CCR-15-1433.

40. Petersen, R.L. (2017). Strategies Using Bio-Layer Interferometry Biosensor Technology for Vaccine Research and Development. Biosensors (Basel) 7. 10.3390/bios7040049.

