## Supplementary materials for "A novel PD-L1-targeted shark V_NAR_ single domain-based CAR-T strategy for treating breast cancer and liver cancer"

#### Cell culture

Human breast cancer cell line MDA-MB-231, human ovarian cancer (OC) cells lines IGROV-1, OVCAR8, and NCI-ADR-RES, human pancreatic cancer (PDAC) cell lines KLM1, Panc-1, and SU8686, and lung cancer cell lines L55, EKVX, and H522 were purchased from American Type Culture Collection (ATCC). MDA-MB-231 was transduced with a lentiviral vector encoding a GFP-firefly-luciferase (GFP-Luc) following our protocol[1]. PD-L1 knockout (KO) MDA-MB-231 cell line was constructed using clustered regularly interspaced short palindromic repeats (CRISPR)/CRISPR-associated protein 9 (CRISPR-Cas9) method. We generated the construct following the design principle as described previously[2]. Briefly, two single-guide RNAs (sgRNAs) targeted endogenous PD-L1 promotor (predicted from the EPD database) were designed and used to subclone into a LentiCRISPRv2 vector and sorted to generate single clones by flow cytometry. The murine melanoma cell line B8979HC (induced in a HGF-tg;CDKN2A$^{fl/fl}$;Tyr-CreERT2-tg mouse by UV and tamoxifen) and the canine tumor cell line Jones was provided by Glenn Merlino (NCI). Hep3B GFP-Luc was established in a previous study[3]. MDA-MB-231 and Hep3B cells were cultured in DMEM supplemented with 10% FBS, 1% L-glutamine, and 1% penicillin–streptomycin; other aforementioned cell lines were cultured in RPMI. Cells were maintained in a humidified atmosphere containing 5% $CO_2$ at 37°C.

#### Phage panning

Phage panning was conducted following our laboratory protocol[4,5]. The Nunc 96-well Maxisorp plate (Thermo Scientific) was coated with 100 µg/ml mPD-L1 in PBS overnight at 4°C. Subsequently, the plate was blocked with 2% bovine serum albumin in PBS for 1 hour at room temperature. Then $10^{10}$-$10^{11}$ CFU of pre-blocked phage supernatant in blocking buffer was added to each well for 1 hour at room temperature to allow binding. After four rounds of panning, the enriched bound phages were eluted with 100 µl pH 2.0 elution buffer at room temperature after four washes with PBS containing 0.05% Tween-20. The eluate was neutralized with 30 µl of 1 M Tris-HCl buffer (pH 8.5) and was used to infect freshly prepared *E. coli* TG1 cells. Single colonies were picked after four rounds of panning, and identified by phage ELISA.

**Antibody production and purification**

The soluble antibody protein-his-flag was produced and purified as previously described[6]. Briefly, the coding sequences of PD-L1 specific $V_{NAR}$s in the pComb3x phagemids were transformed into HB2151 *E.coli* cells. The colonies were pooled for culture in 2 L 2YT media containing 2% glucose, 100 µg/ml ampicillin at 37◦C until the OD600 reached 0.8–1. Culture media was then replaced with 2YT media containing 1mM IPTG (Sigma), 100 µg/ml ampicillin, and shaken at 30°C overnight for soluble protein production. The bacteria pellet was spun down and lysed with polymyxin B (Sigma) for 1 h at 37°C to release the soluble protein. The supernatant was harvested after lysis, and purified using the HisTrap column (Cytiva/GE Healthcare) using AKTA.

**Affinity binding and blocking activity**

All experiments were performed on an Octet instrument (ForteBio) at 30 °C and reagents were prepared in 0.5% BSA, 0.1% Tween20 PBS, pH 7.4 buffer. hPD-L1-His protein was immobilized on to Ni-NTA sensor tips at 5 µg/ml for 120 s. The antigen-coated tips are then dipped into the buffer to stabilize the curve and subsequently dipped into 100 nM or 50 nM B2-hFc or F5-hFc for association and dissociation measurements for a time window of 180 s and 300 s. To detect blocking activity of B2-hFc, hPD-L1-his protein was firstly loaded onto the Ni-NTA sensor tips. Subsequently, the sensor tips were dipped into wells containing 500 nM of $V_{NAR}$-hFc for 300 s, followed by 500 nM hPD-1-hFc protein (Sino Biological) for 300 s, and lastly 300 s of dissociation in the buffer. Raw data was processed using Octet Data Analysis Software 9.0.

**ELISA**

The phage ELISA was performed as previously described[6]. Briefly, the Nunc 96-well Maxisorp plate was coated with 5 μg/ml antigenic proteins, including mPD-L1-his, mPD-L1-hFc, hPD-L1-his, hPD-L1-hFc, and the irrelevant antigen human IgG and PBS in 50 μl PBS per well overnight at 4°C. The plate was blocked with 2% BSA in PBS for 1 hour at room temperature. Pre-blocked phage supernatant was then added to the plate. The binding activity was determined with horseradish peroxidase (HRP)-conjugated mouse anti-M13 antibody (GE Healthcare). To detect whether a shark B2 $V_{NAR}$ is specific to human PD-L1, and not B7-H3, the antigenic proteins hPD-L1 and hB7-H3 were coated onto the 96-well Maxisorp plate. PD-L1 specific binder B2 and B7-H3 specific binder were used to measure the binding affinity and specificity to PD-L1 and B7-H3.

To predict the binding epitope of anti-PD-L1 shark $V_{NAR}$s, we designed total 24 peptides based on the hPD-L1 extracellular domain (ECD) amino acid sequence. Each synthesized peptide (produced by GenScript) was 18 AA in length with 9 AA overlapped, indicating the minimum antigenic region of hPD-L1 ECD recognized by the $V_{NARS}$ can be narrowed down by step-by-step peptide mapping onto a 9-mer peptide epitope. The ELISA was performed in this assay. In brief, a total of 24 peptides were coated onto the 96-well Maxisorp plate. 5ug/ml B12-his-flag, A11-his-flag, and F5-his-flag were then added followed with horseradish peroxidase (HRP)-conjugated anti-flag antibody. The binding activity was determined by OD450. Experiments were performed in triplicate and repeated three times with similar results.

**Flow cytometry**

Surface PD-L1 expression was detected by anti-PD-L1 monoclonal antibody (Biolegend) and goat-anti-human IgG-PE (Jackson ImmunoResearch). Tumor cells were incubated with 10ug/ml of each $V_{NAR}$-his-flag, followed by incubation with mouse anti-flag conjugated with allophycocyanin (APC) (Jackson ImmunoResearch). The transduction efficiency of CAR (B2) T cells was detected by surface anti-EGFR human monoclonal antibody cetuximab (Erbitux) and goat-anti-human IgG conjugated with APC. T cell exhaustion was evaluated via PE PD-1, PE TIM-3, and PE LAG-3 (Thermo Fisher Scientific). 100 μl blood was collected from mice and 1X RBC lysis buffer (eBiosciences) was used to remove red blood cells. To determine the absolute number of CAR T cells in mouse blood, BV711 CD3, Erbitux, and goat-anti-human IgG conjugated with Alexa Fluor 488 CD8 (Biolegend) were used to stain CD3+CAR+ T cells. Counting Beads 123count™ eBeads was used for counting the absolute number of cells. T cell immunophenotyping was performed by surface staining with antibodies against the following antigens: APC-H7 CD4, BV605 CD8, BV421 CD45RA, APC CD62L (BD Bioscience), and PE CD95 (Biolegend). Data acquisition was performed using SONY SA3800 (Sony Biotechnology) and analyzed using FloJo software (Tree Star).

**Western blot**

Cells were lysed with ice-cold lysis buffer (Cell Signaling Technology), and total protein was isolated by centrifugation at 10,000g for 10 minutes at 4°C. Protein concentration was measured using a Bicinchoninic acid assay (Pierce) in accordance with the manufacturer's specifications. For each cell lysates of 20 μg, they were loaded onto a 4-20% SDS-PAGE gel for electrophoresis. Both anti-PDL1 antibody and the anti-GAPDH antibody were obtained from Cell Signaling Technology.

**References**


1. Feng, M., Zhang, J., Anver, M., Hassan, R., and Ho, M. (2011). In vivo imaging of human malignant mesothelioma grown orthotopically in the peritoneal cavity of nude mice. Journal of Cancer *2*, 123-131. 10.7150/jca.2.123.
2. Li, N., Wei, L., Liu, X., Bai, H., Ye, Y., Li, D., Li, N., Baxa, U., Wang, Q., Lv, L., Chen, Y., et al. (2019). A Frizzled-Like Cysteine-Rich Domain in Glypican-3 Mediates Wnt Binding and Regulates Hepatocellular Carcinoma Tumor Growth in Mice. Hepatology *70*, 1231-1245. 10.1002/hep.30646.
3. Li, D., Li, N., Zhang, Y.-F., Fu, H., Feng, M., Schneider, D., Su, L., Wu, X., Zhou, J., Mackay, S., Kramer, J., et al. (2020). Persistent Polyfunctional Chimeric Antigen Receptor T Cells That Target Glypican 3 Eliminate Orthotopic Hepatocellular Carcinomas in Mice. Gastroenterology. https://doi.org/10.1053/j.gastro.2020.02.011.
4. Ho, M., Kreitman, R.J., Onda, M., and Pastan, I. (2005). In vitro antibody evolution targeting germline hot spots to increase activity of an anti-CD22 immunotoxin. J Biol Chem *280*, 607-617. 10.1074/jbc.M409783200.
5. Kim, H., and Ho, M. (2018). Isolation of Antibodies to Heparan Sulfate on Glypicans by Phage Display. Curr Protoc Protein Sci *94*, e66. 10.1002/cpps.66.
6. Feng, M., Bian, H., Wu, X., Fu, T., Fu, Y., Hong, J., Fleming, B.D., Flajnik, M.F., and Ho, M. (2019). Construction and next-generation sequencing analysis of a large phage-

displayed VNAR single-domain antibody library from six naive nurse sharks. Antibody therapeutics *2*, 1-11. 10.1093/abt/tby011.

### Supplementary Figures

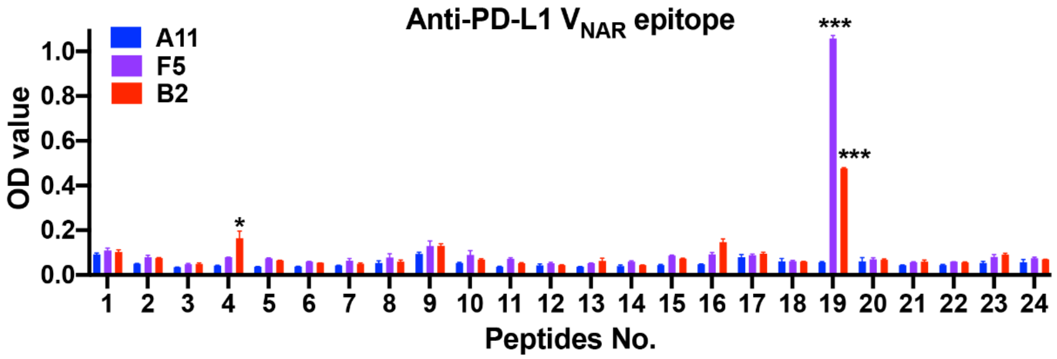


**Supplemental Figure 1.** Epitope mapping of individual anti-PD-L1 $V_{NAR}$s using truncation peptide arrays detected by ELISA. Total 24 peptides were designed based on hPD-L1 ECD. Each peptide is 18 amino acid in length and overlapped 9 AA with adjacent peptide.

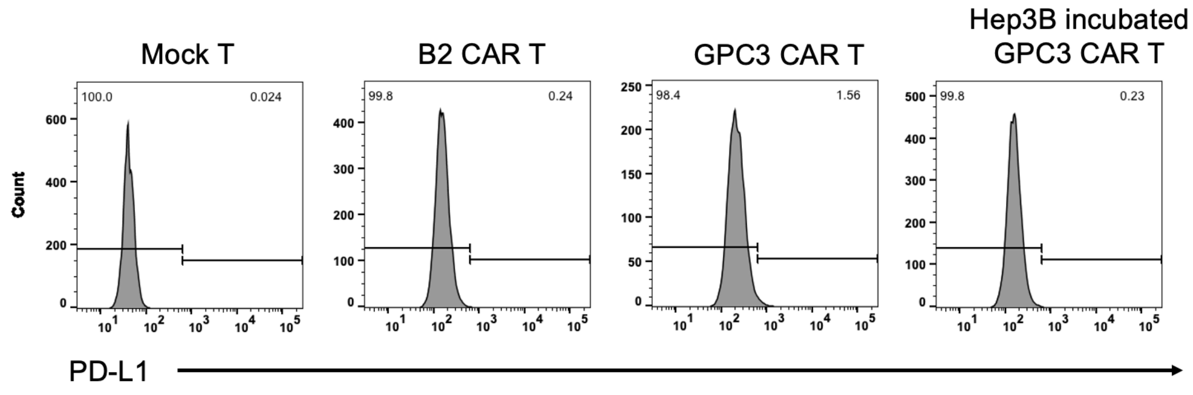


**Supplemental Figure 2.** PD-L1 expression in mock T, CAR (B2) T, GPC3 CAR-T, and Hep3B tumor incubated GPC3 CAR-T cells.

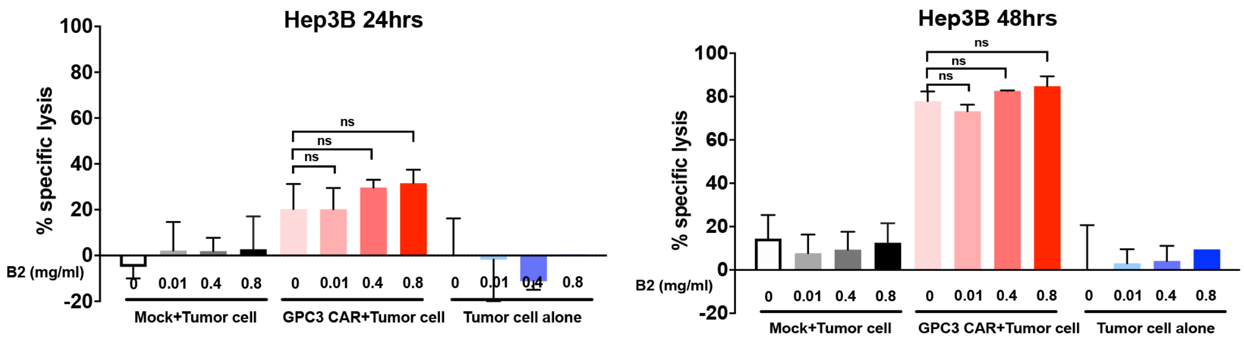


**Supplemental Figure 3.** Monovalent B2 nanobody did not improve GPC3 CAR-T cells killing on Hep3B cells after 24 hours or 48 hours of incubation. Tumor cells alone or mock T cells incubation in the presence of B2 were used as the control in this study. Statistical analyses are shown from three independent experiments. Values represent mean ± SEM. $^{**}P < .01$, $^{***}P < .001$, $^{****}P < .0001$, ns, not significant.
